# Homothallic or heterothallic? A genomic investigation into the sexual capabilities of the ascomycete fungus *Clonostachys rosea*

**DOI:** 10.64898/2026.02.27.708483

**Authors:** David Manyara Wairimu, Andi M. Wilson, Edoardo Piombo, Mikael Brandström Durling, Martin Broberg, Birgit Jensen, Alessandra Ruffino, Sidhant Chaudhary, Henrik H. De Fine Licht, Mukesh Dubey, Magnus Karlsson

**Author notes:** Corresponding author: David Manyara Wairimu, Department of Forest Mycology and Plant Pathology, Swedish University of Agricultural Sciences, Uppsala, Sweden. These authors contributed equally (David Manyara Wairimu, Andi M. Wilson).

## Abstract

Modes of reproduction and sexual strategies strongly influence the genetic diversity and evolutionary potential of a species. The ascomycete fungus *Clonostachys rosea* is reported to be homothallic (sexually self-fertile), although a rapid decay of genome-wide linkage disequilibrium is also reported, something that is not in line with an obligate homothallic mode of reproduction. To investigate this phenomenon, we identified the mating-type (*MAT1*) locus in 66 genome-sequenced *C. rosea* strains under the hypothesis that each strain contains genes from both *MAT1* idiomorphs. Eleven strains indeed contained both *MAT1-1* and *MAT1-2* genes, suggesting homothallism. However, most strains harboured either *MAT1-1* or *MAT1-2* genes and co-existed in North America, Europe and China, suggesting heterothallism. The *MAT1* locus of heterothallic strains was highly conserved, and the linkage disequilibrium half decay distance was 1050 bp, suggesting sexual outcrossing. The presence of conserved *MAT1-1* or *MAT1-2* idiomorphs in strains of other *Clonostachys* species shows that heterothallism is likely the ancestral state. A phylogenetic analysis of 2800 single-copy orthologous genes revealed that homothallic and heterothallic strains separated in two well-supported clades, indicating a single lineage of homothallic *C. rosea*, likely originating in South America, followed by intercontinental dispersal. Homothallic *C. rosea* strains displayed higher nucleotide diversity than heterothallic strains, indicating a lack of outcrossing. This unique case of both homothallic and heterothallic lineages within the same species provides an opportunity to study the genomic consequences of selfing in very closely related strains.

## Introduction

Many fungal species can reproduce both asexually and (Aanen et al., 2023)sexually during their life cycles (Nieuwenhuis & James, 2016). Fungi that are capable of sexual reproduction can typically be described as either heterothallic or homothallic, depending on the precise way they undergo sexual reproduction (reviewed in(Wilson et al., 2021). In heterothallic fungi, mating takes place when there is a physical interaction between two compatible individuals that are of the opposite mating type. In contrast, homothallic fungi can complete the sexual cycle in isolation. The difference in these sexual strategies is usually determined genetically, by genes present at the mating-type (*MAT1*) locus (Lin, 2007).

The *MAT1* locus regulates sexual reproduction in filamentous ascomycetes and controls partner recognition, meiosis, and other mating-related processes. In heterothallic fungi, compatible individuals harbour alternate copies of the *MAT1* locus, termed idiomorphs (Metzenberg & Glass, 1990). Individuals of the MAT1-1 mating type harbour genes associated with the *MAT1-1* idiomorph, while individuals of the MAT1-2 mating type possess *MAT1-2*-associated genes. The completion of sexual reproduction typically requires the expression of both *MAT1-1* and *MAT1-2* genes, hence the requirement that heterothallic species physically interact ahead of sexual development (Min Ni, 2011). Homothallic species, on the other hand, typically harbour both *MAT1-1-* and *MAT1-2-*associated genes within a single genome and/or cell, allowing the expression of both types of genes and thus enabling independent sexual reproduction (Wilson et al., 2015).

Homothallic species tend to evolve from heterothallic ancestors (Yun et al., 1999). This occurs when a genetic event, such as unequal crossing over, combines genes from both *MAT1* idiomorphs within a single genome (Debuchy & Turgeon, 1996). Because of this, the *MAT1* locus structure is often not conserved in homothallic fungi and can differ even amongst close relatives. In some cases, the alternate idiomorphs may even be found unlinked in the genome, resulting in two genomic regions with *MAT1* locus information (Rydholm et al., 2007). For example, self-fertility has evolved independently a number of times in both *Neurospora* (Gioti et al., 2012) and *Aspergillus* (Ojeda-López, 2018) with individual homothallic species harbouring different *MAT1* locus structures depending on the precise mechanisms that brought the *MAT1* genes together.

The *MAT1* genes can be classified as primary and secondary genes (Wilson et al., 2021). The primary *MAT1* genes are the defining genes of each idiomorph and are almost universally present at the relevant idiomorph. The primary *MAT1* genes are *MAT1-1-1* and *MAT1-2-1*, both of which encode transcription factors (Ferreira et al., 1998; Klix et al., 2010). The MAT1-1-1 protein harbours an alpha-box domain, and the MAT1-2-1 protein harbours an HMG-box. Both genes have been functionally characterized in a variety of model and non-model fungi and have proven to be essential for sexual development in almost all cases (Coppin et al., 1997; Ferreira et al., 1998; Glass et al., 1990; Klix et al., 2010). They both contribute to the initial formation of reproductive tissue, regulation of the pheromone system, and production of ascomata. The secondary *MAT1* genes (e.g. *MAT1-1-2, MAT1-1-3*) are less conserved and typically show lineage-specific presence (Wilken et al., 2017). There is a paucity in functional data for these genes, and where this data does exist, there are species-specific differences in their functions. For example, the *MAT1-1-3* gene is entirely dispensable for sexual reproduction in *Sordaria macrospora* (Klix et al., 2010) but essential for fertility in *Fusarium graminearum* (Zheng et al., 2013).

The lack of outcrossing in obligate homothallic species is predicted to result in the accumulation of deleterious mutations, a process known as Muller’s ratchet (Felsenstein, 1974; Muller, 1932, 1964). Over the long term, this continuous accumulation of harmful mutations can reduce fitness and increase extinction risk, especially in small or strictly selfing populations (Gioti et al., 2013; Loewe & Cutter, 2008; Pamilo et al., 1987). Fungal systems with suppressed recombination around the *MAT1* locus show signatures consistent with relaxed selective constraint and mutation accumulation, including homothallic *Neurospora* species and anther-smut fungi *Microbotryum*, where non-recombining mating-type regions exhibit degeneration such as frameshifts and premature stop codons in *MAT1* genes (Fontanillas et al., 2015; Wik et al., 2008). In the short term, however, obligate homothallism may allow the conservation of favourable combinations of alleles, providing high fitness to a homothallic, essentially clonal, lineage. Experimental work in the homothallic fungus *A. nidulans* has shown that selfed progeny can exhibit higher mean fitness than outcrossed progeny, despite reduced recombination (López-Villavicencio et al., 2013), and similarly successful clonal or selfing lineages have been described among plant-associated pathogens and mutualists, including *Phytophthora infestans* and *Epichloe* species (Drenth et al., 2019; McDonald & Linde, 2002).

The fungus *Clonostachys rosea* (Schroers, 1999) is a filamentous ascomycete from the *Hypocreales* order. It is the anamorph of the teleomorph *Bionectria ochroleuca* (Schwein.) (Schroers, 1997, 1999; Schroers, 2001). *Clonostachys rosea* is the recommended species name under the one fungus, one name principle (Rossman, 2014). Two variants of *C. rosea* can be found in the literature, *C. rosea* forma (f.) *rosea* and *C. rosea* f. *catenulata*, although they constitute a single species based on genealogical concordance phylogenetic species recognition (Moreira et al., 2016). The sexual fruiting body, the perithecium, is mostly found in pantropical and subtropical regions, while conidial (anamorph) strains of *C. rosea* are cosmopolitan (Schroers, 1999). *Clonostachys rosea* was reported to be homothallic, as strains originating from ascospores frequently produce perithecia in single cultures on rich media (Schroers, 1999). Notably, conidial strains only rarely form perithecia under the same conditions. However, a population genomic study of *C. rosea* (Broberg et al., 2018) reported rapid decay of genome-wide linkage disequilibrium (LD), which is not expected given an obligate homothallic mode of reproduction.

Strains of *C. rosea* are typically isolated from a wide variety of ecological niches, including soil, other ascomycete fungi, plant litter and from living plants (García, 2003; Mueller, 1986; Nobre, 2005; Walker, 1975), but rare isolations from nematodes and arthropods are also reported (Haarith, 2020; Verdejo-Lucas, 2002). This broad habitat distribution of *C. rosea* can be correlated with an ecological generalist lifestyle, which includes plant endophytism, rhizosphere competence, polyphagous ability and mycoparasitism (Chatterton, 2012; Li, 2002; Maillard, 2020; Saraiva, 2015; Shigo, 1958). The broad nutritional versatility that characterises generalist behaviour in *C. rosea* is suggested to be important for the ability of certain strains to control plant diseases and their commercial use as biological control agents (Jensen, 2022).

To shed light on the reproductive mode of *C. rosea*, we performed genome-sequencing and identified the *MAT1* locus in 66 *C. rosea* strains under the hypothesis that each strain contains genes from both *MAT1* idiomorphs, as expected from the reported homothallism. Although eleven strains did indeed contain both *MAT1-1* and *MAT1-2* genes, most strains harboured either *MAT1-1* or *MAT1-2* genes, suggesting heterothallism. A phylogenetic analysis of 2800 single-copy orthologues revealed that homothallic and heterothallic strains clustered in two separate clades, indicating a single lineage of homothallic *C. rosea*, likely originating in South America, followed by inter-continental dispersal. The presence of both homothallic and heterothallic lineages within a single species provides an excellent opportunity to study the genomic consequences of selfing.

## Materials and methods

### Fungal strains, culture conditions and DNA extraction

*Clonostachys rosea* strains (Table 1 in Supplemental File S1) were revived from glycerol stocks stored at -70°C at the Department of Forest Mycology and Plant Pathology at the Swedish University of Agricultural Sciences, and maintained on potato dextrose agar (PDA, Oxoid, Cambridge, UK) at 25°C in darkness. For DNA extraction, strains were grown in 200 ml liquid Czapek-Dox medium (Sigma-Aldrich, Steinheim, Germany), Vogel’s minimal medium (Vogel, 1956) or malt extract (1.75 %) with peptone (0.25 %) medium at room temperature, shaking at 120 rpm. Cultures were harvested after 3-13 days, depending on growth rate, by snap freezing in liquid nitrogen, and then freeze-dried. High-quality genomic DNA was extracted using a cetyltrimethylammonium bromide (CTAB)/chloroform-based protocol or the Qiagen-tip 100 kit (Qiagen, Hilden, Germany) as described previously (Broberg et al., 2018).

### Genome sequencing and assembly

Base coverage for ten *C. rosea* genomes was generated using Illumina HiSeqX paired-end sequencing with a 350 bp insert size and 150 bp read length following standard library preparation kits. Sequence read data for these ten *C. rosea* strains, together with reads from 52 previously sequenced *C. rosea* strains (Broberg et al., 2018), were assembled with ABySS ver. 1.3.6 (Simpson et al., 2009), as described by Broberg et al. (2018). Assembly completeness was assessed with Benchmarking Universal Single-Copy Orthologs (BUSCO) v.5.1.2 (Seppey et al., 2019) using the “sordariomycetes odb12” lineage dataset, and scaffolds shorter than 500 bases were removed using the “funannotate sort” function in Funannotate v.1.8.15 (Palmer & Stajich, 2020).

### Genome annotation and gene prediction

RNA-seq data from *C. rosea* strains IK726, ACM941, and 88-710 (Table 2 in Supplemental File S1) were retrieved and adapter- and quality-trimmed using bbduk v.38.90 (Bushnell et al., 2014). For samples generated from *C. rosea* grown in interaction with other organisms (e.g. plants or plant pathogens), trimmed reads were mapped to the *C. rosea* IK726 ver. 2 reference genome (GCA_902827195.2) (Broberg et al., 2018) using STAR v.2.7.9a (Dobin et al., 2013) with default parameters. Reads that did not align to the *C. rosea* genome were identified with “samtools view” v.1.19 (Danecek et al., 2021) and removed using “seqtk subseq” v.1.2-r101 (Li, 2021). Transcriptomes were then assembled with Trinity v.2.11.0 (Grabherr et al., 2011) using the option “--jaccard_clip”.

Funannotate v.1.8.15 (Palmer & Stajich, 2020) was used to filter and rename the scaffolds of the 62 newly assembled *C. rosea* genomes, together with four additional *C. rosea* genomes (strains ACM941, CanS41, IK726 and 88-710), four *C. farinosa* genomes (synonym *C. byssicola*), four *C. chloroleuca* genomes, three *C. rhizophaga* genomes, one *C. solani* genome, and one *C.* sp. CBS 192.96 genome retrieved from the National Center for Biotechnology Information (NCBI) and Mycocosm (Table 3 in Supplemental File S1). All genomes underwent repeat identification with RepeatModeler v.2.0.1 (using the “LTRStruct” option) and repeat masking with RepeatMasker v.4.1.1 using default parameters (Flynn et al., 2020). Funannotate was trained for *C. rosea* gene prediction using the *C. rosea* IK726 ver. 2 reference genome (GCA_902827195.2) (Broberg et al., 2018) and the options “--optimize_augustus --organism fungus –busco_db sordariomycetes”, incorporating Trinity-assembled transcripts as transcript evidence and UniProt protein sequences obtained via “funannotate setup” as protein evidence. The trained gene model was then applied to predict genes in each newly assembled *C. rosea* genome using the same parameters except for “optimize_augustus”. Functional annotation of predicted proteins was performed with InterProScan v.5.48-83.0 (Jones et al., 2014) using the options “--iprlookup --goterms --pathways”. Secretome prediction followed the pipeline of Piombo et al. (2023). These annotations were integrated using “funannotate annotate” v.1.8.15 (Palmer & Stajich, 2020), with the options “--signalp --iprscan --busco_db sordariomycetes”.

Regarding nomenclature, we here use uppercase letters in italics for the *MAT1* locus and the *MAT-1-1*/*MAT1-2* idiomorphs (Turgeon & Yoder, 2000), while we use lowercase letters in italics for genes in *Clonostachys* (Jensen et al., 2022). The *MAT1* locus and corresponding *mat* genes from strains *C. rosea* CBS 100502 (*MAT1-1*) and CBS 103.94 (*MAT1-2*) were identified using the MAT1-1-1 and MAT1-2-1 proteins from *E. festucae* (NCBI accession numbers AKH05039.1 and AEI72619.1, respectively) and a local tBLASTn analysis. Up to ten predicted proteins upstream and downstream of *mat1-1-1* and *mat1-2-1* were used in BLASTp similarity analyses against the NCBI nr database to determine the extent of the *MAT1* locus in these two strains. In both idiomorphs, the *MAT1* locus was flanked by homologs of APN2 and SLA2. These full length *MAT1-1* and *MAT1-2* loci were subsequently used in local BLASTn analyses against the remaining 77 genomes to determine the *mat* gene content and predict the mating type. CLINKER as implemented in CAGECAT (Gilchrist & Chooi, 2021) was used to generate synteny maps of the *MAT1* loci of selected strains. Mating type and sexual strategy was mapped onto a world map using mapPies in the R package rworldmap (South, 2011).

### Phylogenomic analyses

The BUSCO proteins common to all 66 *C. rosea* genomes as well as the 13 additional *Clonostachys* species were aligned using MUSCLE ver. 5.1.0 (Edgar, 2004) with default settings. The subsequent alignments were trimmed using trimAl ver. 1.1.4 (Capella-Gutiérrez et al., 2009) and the –automated1 flag. Each alignment was used to infer single-protein maximum likelihood trees using IQTree ver. 2.2.2.7 (Minh et al., 2020; Nguyen et al., 2015) with 1000 ultrafast replicates for bootstrapping (-B 1000). A combined tree was then estimated using ASTRAL ver. 5.7.7 (Zhang et al., 2018), using local posterior probabilities (LPP) as a measure of branch and topological support. An LPP value close to 1 suggests high support for that node and that there is congruence between the individual gene trees for that node. The final tree was visualized in FigTree ver. 1.4.4. (http://tree.bio.ed.ac.uk/software/tree/) and edited in Affinity Designer 2.

### Read alignment, SNP calling, and quality filtering

Illumina paired-end reads from 63 *C. rosea* strains (Table 1 in Supplemental File S1), including *C. rosea* IK726 ver. 1 (Karlsson, 2015) were adapter- and quality trimmed using bbduk v.38.90 (Bushnell et al., 2014) with default settings. The success of the trimming was determined with fastQC v.0.11.9 (https://www.bioinformatics.babraham.ac.uk/projects/fastqc) and cleaned reads were mapped to the *C. rosea* IK726 ver. 2 genome (GCA_902827195.2) (Broberg et al., 2018) using Bowtie2 v.2.4.2 (Langmead & Salzberg, 2012). Duplicate reads were identified with Picard MarkDuplicates v.2.18.29 (https://broadinstitute.github.io/picard) and BAM files with flagged duplicates were indexed using SAMtools v.1.19 (Danecek et al., 2021). Single nucleotide polymorphisms (SNPs) between the mapped reads and the *C. rosea* IK726 ver. 2 reference genome were called with BCFtools v.1.12 (Danecek & McCarthy, 2017) using the parameters “-excl-flags UNMAP,SECONDARY,QCFAIL,DUP” for “bcftools mpileup” and “-c --ploidy 1 -v -Oz” for “bcftools call”. Initial filtering of the resulting Variant Call Format (VCF) file was performed with VCFtools v.0.1.17 (Danecek et al., 2011) using the following criteria: “--max-missing 0.25, --min-meanDP 40, --minQ 500, --minDP 10, --maf 0.05, --remove-indels, and --recode”.

To minimize SNP detection errors, further quality filtering retained only bi-allelic SNPs with sequencing depth between 55 and 92 (mean genome wide coverage 73.61). Based on the structure of the *MAT1* locus, the resulting dataset was then partitioned into homothallic and heterothallic strains for downstream analyses. Additional filtering was applied separately to each group: loci were required to have at least 20 reads, and strong allelic support within each strain was required, such that the called allele (reference or alternate) was supported by at least 90% of reads at that site. Loci not meeting these criteria were coded as missing data. Finally, SNPs with more than 25% missing data and a minor allele frequency (MAF) below 5% were removed from each dataset.

### Population structure analyses

Genetic structure among the 63 *C. rosea* strains (Table 1 in Supplemental File S1) was analysed using principal component analysis (PCA) implemented in the R package SNPRelate v.1.4.0 (Zheng et al., 2012), and model-based clustering in LEA v.3.18.0 (Frichot & François, 2015). The analyses were performed on the original VCF file containing SNPs of all 63 strains, generated after SNP calling but prior to quality filtering. To reduce LD, SNPs were pruned with PLINK v.1.90b7 (Purcell et al., 2007) using a window size of 50 SNPs, a step size of 5 SNPs, and an LD threshold of 0.5. The dataset was further filtered to retain SNPs with coverage > 20, no missing data, and MAF > 10%. PCA was performed using the function snpgdsPCA in R v.4.4.2 (R Core Team 2025), and the first two eigenvectors were plotted to visualize population structure. Population structure was further assessed using sparse nonnegative matrix factorization (sNMF) (Frichot et al., 2014), which was used to estimate ancestry proportions and admixture levels for each strain. Twenty independent runs were carried out for *K* = 1–10, and the minimum cross-entropy was observed at *K* = 6.

A phylogenetic network was also constructed to investigate reticulate evolutionary relationships among the 63 strains. For this purpose, the LD-pruned and filtered VCF file was converted to NEXUS format using the vcf2phylip v.2.0 script (Ortiz, 2019), and a neighbour-joining phylogenetic network was then inferred in SplitsTree v.4.19.2 (Huson & Bryant, 2006).

### Population genetic analyses

Filtered SNP datasets for homothallic and heterothallic strains, generated as described in the previous section, were used for all population genetic analyses. Minor allele frequency spectra, Tajima’s D, and nucleotide diversity (π) metrics were used to assess and quantify levels of variation between homothallic and heterothallic groups. Minor allele frequencies were calculated using the “--freq” option in VCFtools v.0.1.17 (Danecek et al., 2011) and visualized with ggplot2 in R v.4.4.2 (R Core Team 2025). Tajima’s D (Tajima, 1989) and nucleotide diversity (Tajima, 1983) were calculated separately for homothallic and heterothallic datasets in 10 kb sliding windows with 5 kb steps using the vcfR package v.1.14.0 (Knaus & Grünwald, 2017). Differences in Tajima’s D and nucleotide diversity between groups were assessed via 1000 bootstrap resamplings within each group, with mean values recalculated per iteration and compared using 95% confidence intervals and empirical *P*-values (significant if *P* < 0.05 or *P* > 0.95).

Linkage disequilibrium (LD) was quantified from pairwise r^2^ values (Hill & Robertson, 1968) between SNPs using VCFtools v.0.1.17 (Danecek et al., 2011) with the “--haploid” flag. Pairwise r^2^ values were calculated within 10 kb windows (5 kb step size), and mean r^2^ per window was used to assess LD decay. Physical distances between SNPs were binned into 100 bp intervals, and mean r^2^ per bin was modelled with LOESS smoothing in R. The distance at which LD decayed to half its maximum was estimated as the midpoint of the bin where the LOESS curve crossed 50% of the maximum r^2^. Ninety-five percent confidence intervals were obtained by 1000 bootstrap resampling, and differences between homothallic and heterothallic groups were tested using a Wilcoxon rank-sum test.

Shared and group-specific SNPs were identified using BCFtools v.1.12 (Danecek & McCarthy, 2017), where shared SNPs segregated in both groups and group-specific SNPs occurred in only one group. Minor allele frequencies for each SNP set were calculated using VCFtools v.0.1.17 and visualized with ggplot2. To test for differences in nonsynonymous mutation accumulation associated with sexual strategy, nucleotide diversity was calculated separately for the predicted nonsense, missense, and synonymous SNPs in shared and group-specific datasets. SNP functional impacts were predicted and annotated using SnpEff v.5.2e (Cingolani et al., 2012): predicted high-impact (nonsense) SNPs are defined as non-synonymous changes predicted to disrupt protein function via stop codon gain or loss; predicted moderate-impact (missense) SNPs are non-synonymous changes potentially altering protein sequence without disrupting function; and predicted low-impact (silent) SNPs are generally synonymous and neutral. Statistical significance of nucleotide diversity differences was evaluated using 1000 bootstrap resamplings, with empirical *P*-values defined as the proportion of iterations in which nucleotide diversity from one group exceeded that of the other (significant if *P* < 0.05 or *P* > 0.95).

To further examine selection pressures between homothallic and heterothallic groups, pN/pS ratios were calculated using all SNPs segregating within each group for homothallic and heterothallic groups. pN/pS ratios were calculated as the number of nonsynonymous SNPs per nonsynonymous site divided by the number of synonymous SNPs per synonymous site. Ratios above one indicates positive selection favouring protein changes, ratios below one suggest purifying selection against deleterious changes, and a ratio of one implies neutrality (Hurst, 2002; Tanaka & Nei, 1989). Ratios were computed for each gene with SNPs using a custom Python script (https://github.com/markhilt/mutation_analysis/blob/main/dnds.py), which counts potential synonymous and nonsynonymous sites per gene based on the reference, and the observed synonymous and nonsynonymous SNPs, assuming closely related strains and therefore no correction for multiple substitutions. Gene-wise pN/pS ratios were then averaged within homothallic and heterothallic groups, and differences in selection levels between groups were tested using Welch’s t-test and Wilcoxon rank-sum test in R v.4.4.2 (R Core Team 2025).

### Phenotypic analyses

The ability of homothallic strains to produce perithecia in single culture was investigated by inoculating strains on PDA (Oxoid, Cambridge, UK) in Petri dishes in duplicates, incubation at 20°C, followed by daily examination for one month using an Olympus SZD-ILLD dissection microscope to observe mature perithecia. Release of asci and ascospores from perithecia were observed using an Olympos BX 60F5 microscope. Images were captured with a Canon DS 124491 camera. The ability of perithecia formation in crosses between selected *MAT1-1* and *MAT1-2* heterothallic strains was investigated by dual confrontation assays on PDA and synthetic crossing (SC) medium (Westergaard & Mitchell, 1947) and incubation at 20°C for one month. Three biological replicates for each medium were included. Furthermore, dual confrontation assays between selected homothallic strains were performed in duplicates on PDA and incubated at 20°C to assess self/non-self-recognition and hyphal fusion.

## Results

### Features of the *Clonostachys rosea* genomes

The 62 *de novo* short-read Illumina genome assemblies generated in this study had an average size of 57.96 Mb, ranging from 53.39 Mb (*C. rosea* CBS 193.94) to 62.27 Mb (*C. rosea* 1829). The predicted number of protein-coding genes averaged 18586 per genome, with a range of 17638 (*C. rosea* SHW-3-1) to 19366 (*C. rosea* 1829). Assemblies contained an average of 1133 scaffolds (range: 227 for *C. rosea* SHW-3-1 to 2369 for *C. rosea* CBS 649.80).

Genome completeness assessed using the “sordariomycetes odb12” BUSCO lineage dataset ranged from 88.3% to 98.7%, indicating near-complete recovery of conserved genes (Table 4 in Supplemental File S1). A total of 2805 complete predicted BUSCO proteins were shared among all 79 *Clonostachys* genomes that were included in this study. Out of these, five were excluded as phylogenetically uninformative, leaving 2800 genes for downstream phylogenomic analyses.

### Distribution of mating-type genes in *Clonostachys* strains

Based on sequence similarity analyses, *mat* genes were identified in all 79 *Clonostachys* genomes, typically found within complete *MAT1* loci and flanked by the *apn2* and *sla2* genes (Figure 1A-B, Table 5 in Supplemental File S1). Nineteen of the *C. rosea* strains harboured typical *MAT1-1* loci, including the *mat1-1-1*, *mat1-1-2*, and *mat1-1-3* genes. A further 36 *C. rosea* strains harboured typical *MAT1-2* loci, possessing the *mat1-2-1* gene as well what appears to be a homolog of the *Fusarium mat1-2-9* gene. Additional small genes were also identified in several of the *MAT1* loci, some of them with similarity in both *MAT1-1* and *MAT1-2* idiomorphs (Figure 1A). These were considered annotation artefacts given their presence/absence patterns, presence within both idiomorphs, and short lengths. The *mat* genes found in *C. rosea* were also found in *C. chloroleuca*, *C. farinosa*, *C. rhizophaga*, *C. solani*, and *C.* sp. CBS 192.96, always in separate *MAT1-1* or *MAT1-2* idiomorphs in each strain (Figure 1B, Table 5 in Supplemental File S1). Presence of either the *MAT1-1* or *MAT1-2* idiomorph in these strains suggests heterothallism in most *Clonostachys* species.

**Figure 1.**
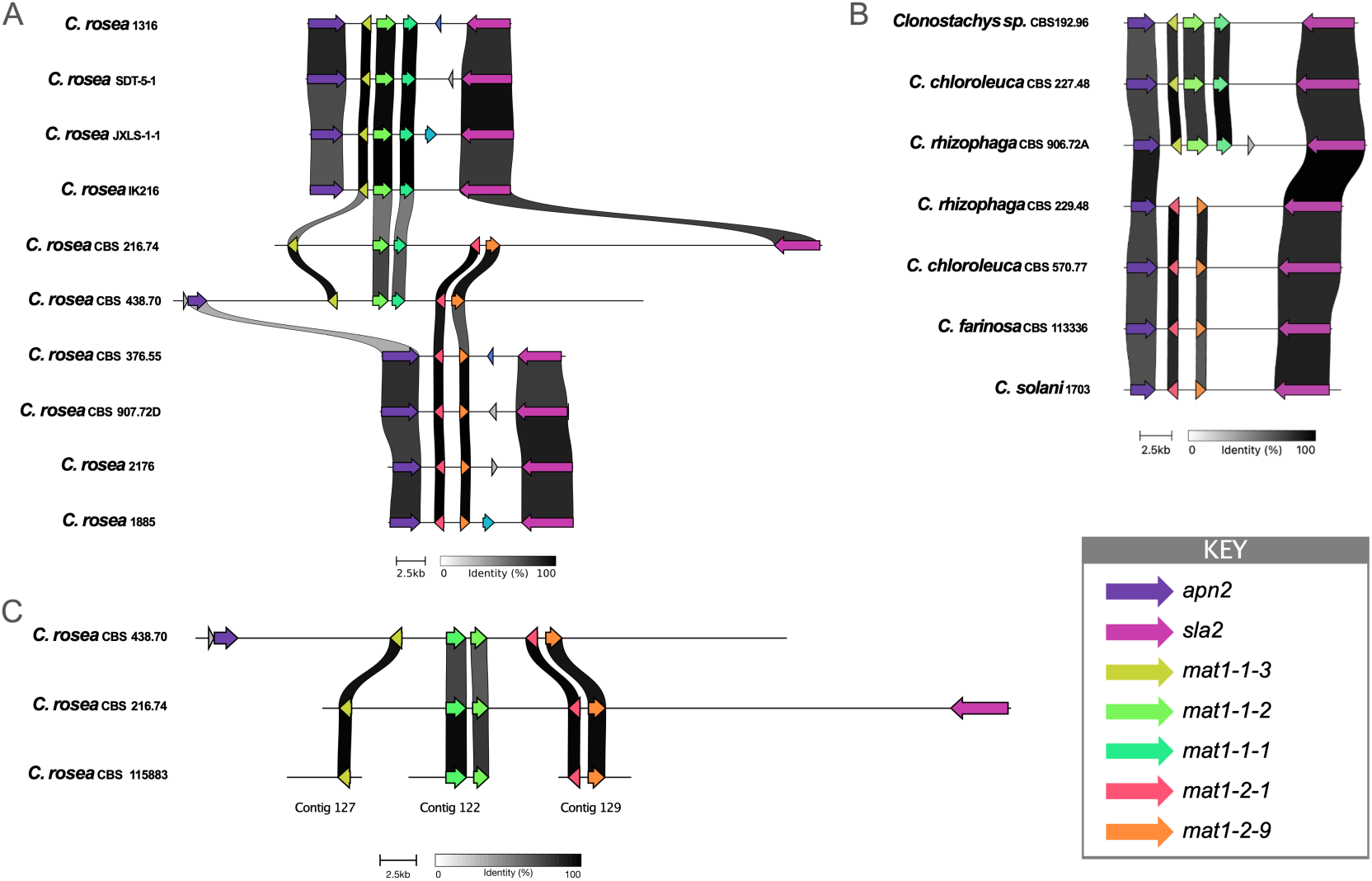
*MAT1* locus organization in *Clonostachys*. **(A)** Synteny maps of the *MAT1* loci in selected heterothallic and homothallic *C. rosea* strains. The *MAT1* locus in *C. rosea* is flanked by *apn2* and *sla2,* a typical arrangement in other Pezizomycotina species. In heterothallic strains, the *MAT1-1* idiomorph harbours the *mat1-1-1*, *mat1-1-2* and *mat1-1-3* genes, while the *MAT1-2* idiomorph harbours the *mat1-2-1* and *mat1-2-9* genes. In homothallic strains, all five *mat* genes are found in an expanded locus, separated by highly repetitive sequence. Synteny maps of the *MAT1* loci in selected *Clonostachys* species. The *MAT1* locus structure supports heterothallism in all species, where only either the *MAT1-1* or *MAT1-2* idiomorph is present per strain. For both *C. rhizophaga* and *C. chloroleuca*, both *MAT1-1* and *MAT1-2* strains were present, further supporting heterothallism as the sexual strategy in these species. **(C)** Synteny maps of contigs harbouring *mat* genes in selected homothallic *C. rosea* strains. Only two of the *C. rosea* homothallic strains harboured almost fully intact loci, while the *mat* genes for the remaining nine homothallic strains (represented here by strain CBS 115883) were scattered throughout the genome assembly on much smaller contigs, likely due to assembly fragmentation caused by repetitive sequences within the expanded locus. In all panels, gene orientation is indicated by arrow direction; grey arrows denote putative annotation artifacts inferred from inconsistent presence/absence patterns and occurrence in both *MAT1-1* and *MAT1-2* idiomorphs.

The remaining eleven *C. rosea* strains possessed all five *mat* genes, representing both the *MAT1-1* and *MAT1-2* idiomorphs, suggesting homothallism. Interestingly, these genes were found in greatly expanded *MAT1* loci of more than 40 kb. In contrast, the *MAT1-1* and *MAT1-2* idiomorphs from the heterothallic strains ranged from 12 to 18 kb in length, from *apn2* to *sla2* (Figure 1A). Notably, of the eleven genomes from homothallic strains, only CBS 216.74 and CBS 438.70 had almost fully assembled *MAT1* loci, with all five *mat* genes on a single contig. In the other nine homothallic strains, the genes were present on much shorter contigs, typically with the *mat1-1-1* and *mat1-1-2* genes on a single contig, *mat1-1-3* on a single contig, and *mat1-2-1* and *mat1-2-9* on a single contig (Figure 1C). This is likely the result of the *mat* genes in these strains being interspersed by AT-rich, highly repetitive DNA. This sequence type is typically very difficult to assemble into contiguous sequence.

After one month of growth on PDA, there were varying amounts of perithecia, asci and ascospores produced by the homothallic strains (Figure 2). When homothallic strains were confronted with each other in dual cultures, there was no production of perithecia in the interaction zones after one month, showing that perithecia production was not triggered by non-self-interaction. Typically, interactions either consisted of a clearing zone between the strains or profuse production of aerial mycelium with no clear sign of hyphal anastomosis. No perithecia was observed in single cultures of any heterothallic strain. Dual confrontation of selected heterothallic strains carrying opposite *MAT1* idiomorphs did not result in production of perithecia on PDA or SC medium after one month.

**Figure 2.**
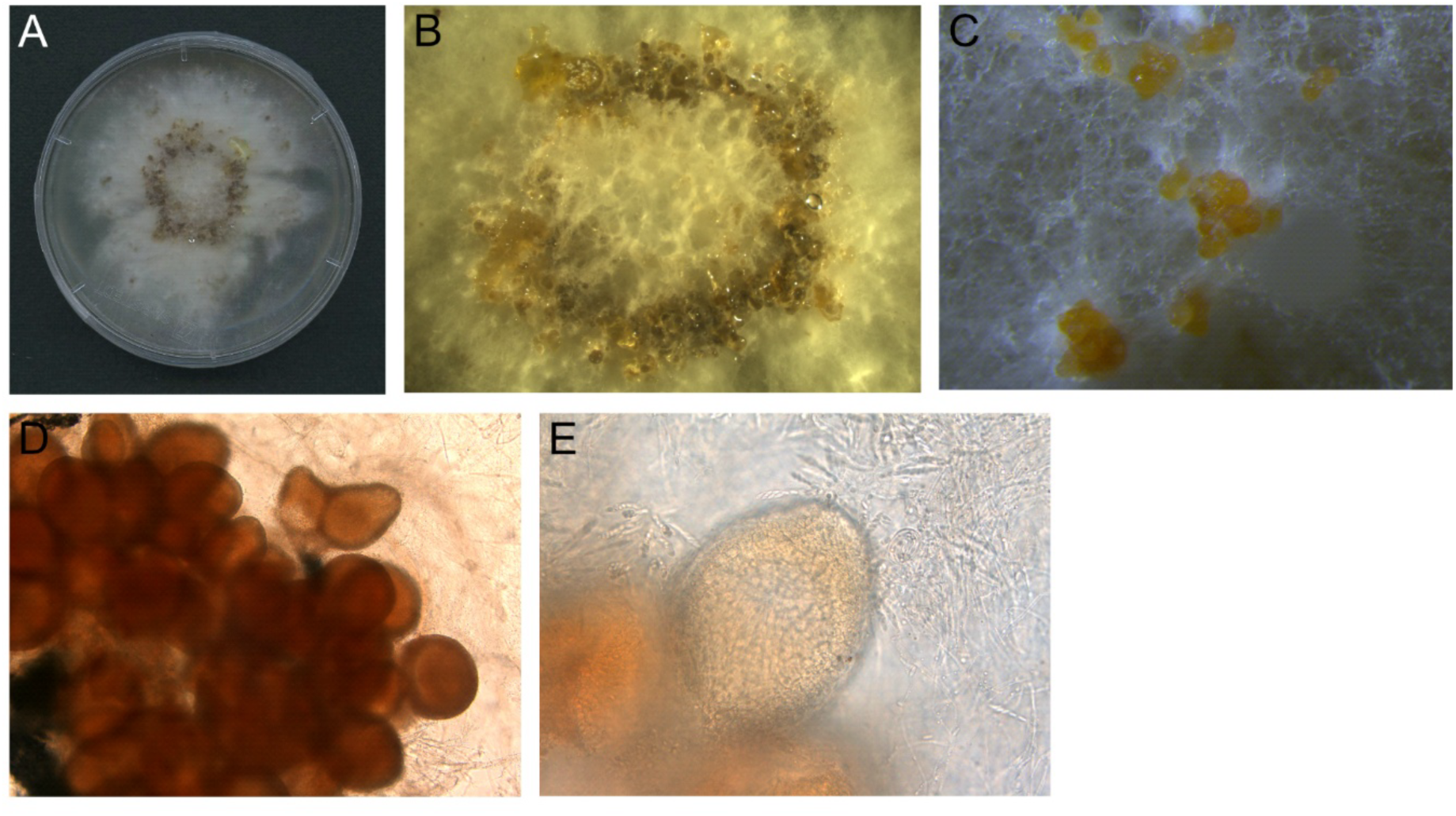
Production of perithecia in homothallic *C. rosea* strains. Homothallic *C. rosea* strains were grown on PDA and incubated at 20°C. After two weeks, perithecium formation was visible and examined daily for the following 2 weeks. (**A, B**) Macroscopic view of perithecia formation in *C. rosea* strain CBS 115883. (**C, D**) Perithecia formation in *C. rosea* strain B15. (**E**) Release of asci from perithecia of *C. rosea* strain CBS 289.78.

### Phylogenetic relationships of *Clonostachys* strains

The phylogenetic analysis of 2800 BUSCO genes clustered heterothallic and homothallic strains into two separate lineages with an LPP value of 1, strongly indicating that these two lineages have diverged (Figure 3). Within the homothallic lineage, one group contained two strains from New Zealand (B14 and B15), one strain from Argentina (CBS 115883) and one strain from Chile (CBS 222.93). Among the remaining strains, SHW-3-1 from China and CBS 438.70 from Japan clustered together, while the remaining strains originated from Brazil, Jamaica, Mexico and Venezuela. Among heterothallic strains, some clustering according to geographic origin was evident, as most strains from China and some strains from Slovenia clustered in distinct groups. Mapping mating types and/or sexual strategy geographically showed that heterothallic strains containing *MAT1-1* or *MAT1-2* mating types existed within the same areas in North America, Europe and China (Figure 4), suggesting the opportunity for sexual reproduction. Countries with single mating types were typically only represented by a single strain and likely represent a sampling bias rather than a true mating type bias. The homothallic strains were restricted to South and Central America, Japan, China and New Zealand, further supporting the single origin of this lineage and a single origin of homothallism in this species.

**Figure 3.**
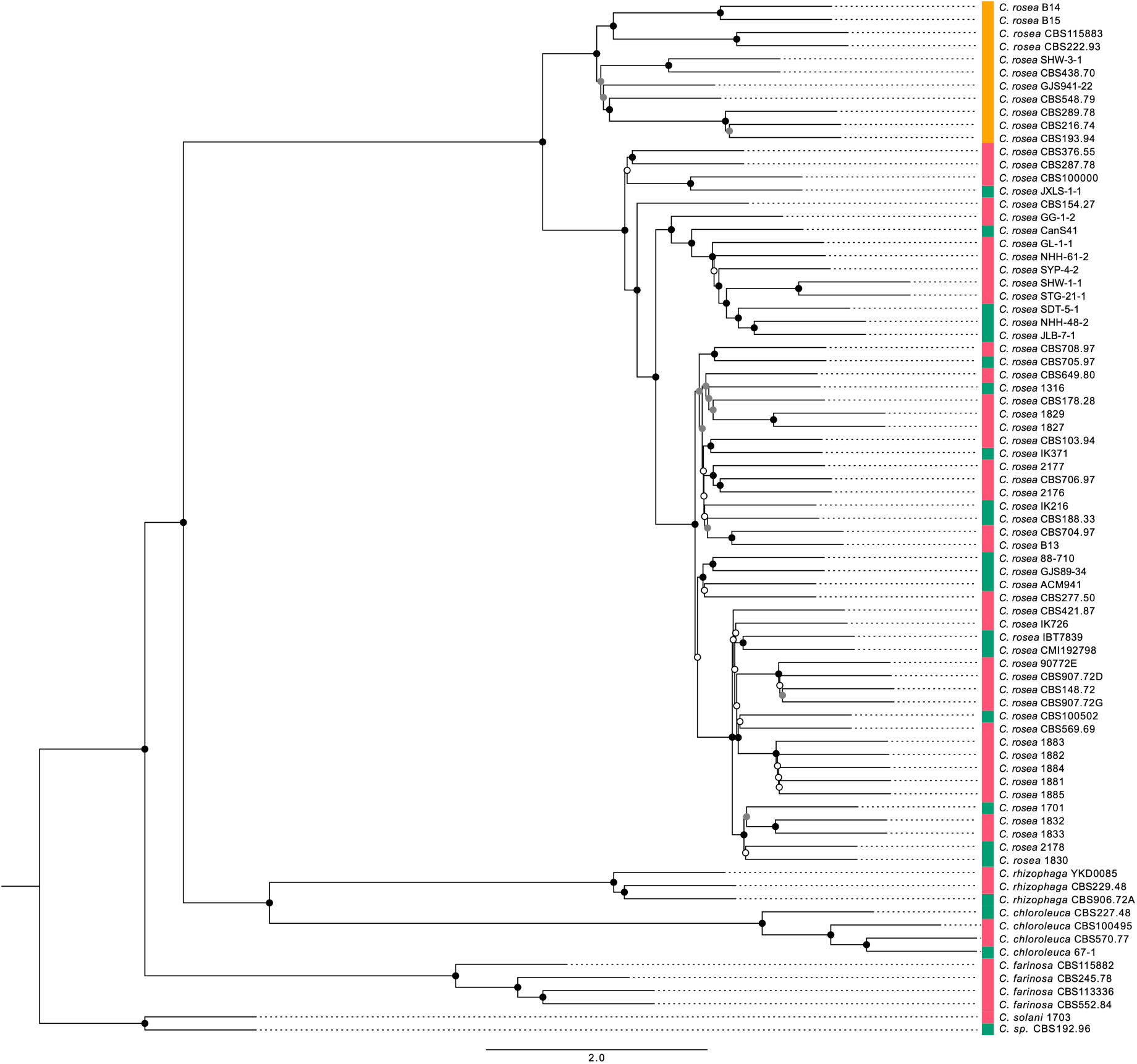
The phylogenetic relationship between homothallic and heterothallic strains reveals a single origin of homothallism in *C. rosea.* The phylogeny is based on 2800 BUSCO proteins present in the genomes of all 66 *C. rosea* strains and additional outgroups. Node support, measured by local posterior probabilities (LLP), are illustrated by black (LPP = 1), grey 0.80 < LPP < 0.99), and white (LPP < 0.8) circles positioned on each node. The track next to each strain represents sexual strategy. Strains harbouring both *MAT1-1* and *MAT1-2* idiomorphs, indicating homothallism, annotated in orange. Strains harbouring the *MAT1-1* idiomorph (annotated in green) or the *MAT1-2* idiomorph (annotated in pink) are likely heterothallic. This analysis supports the clustering of the homothallic strains separately from the heterothallic strains.

**Figure 4.**
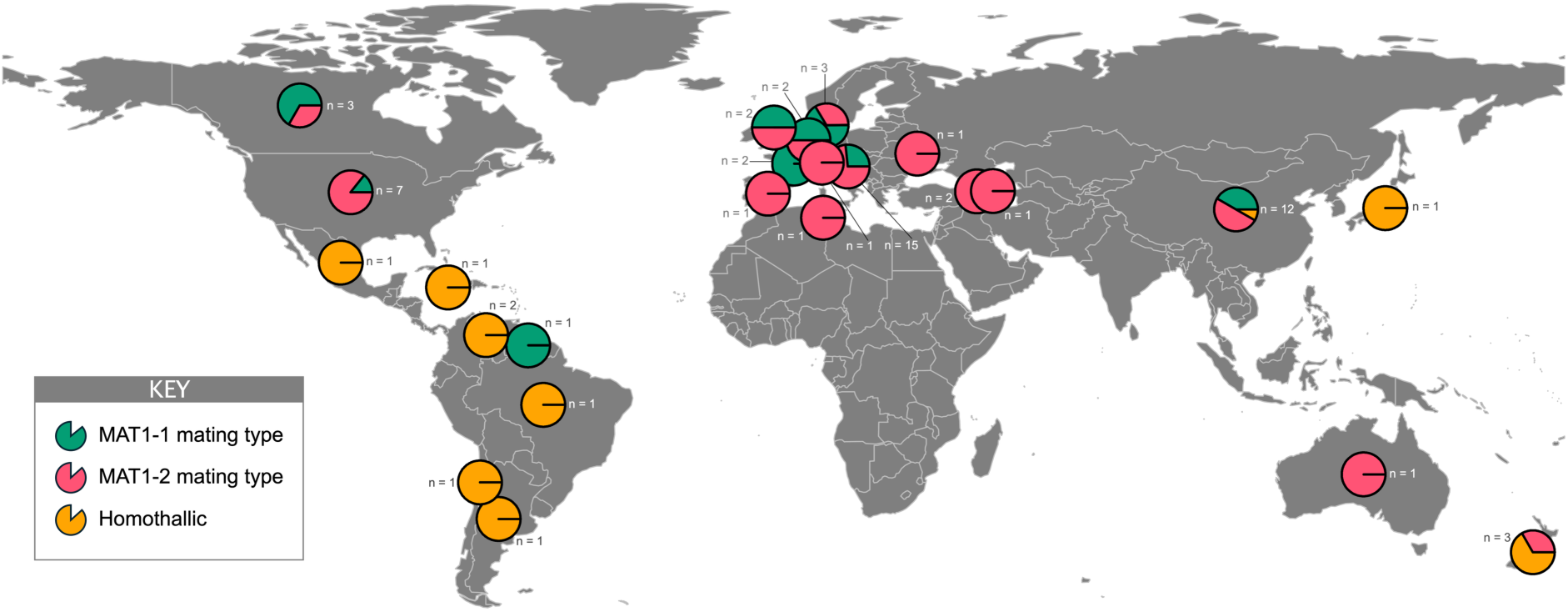
Geographic distribution of heterothallic and homothallic *C. rosea* strains. Homothallic strains, coloured in orange, mainly originate from South and Central America, while heterothallic strains, coloured in green and pink for the *MAT1-1* and *MAT1-2* mating types, respectively, mainly originate from North America, Europe and Asia. This supports the hypothesis of a single origin of homothallism, likely in South America, and subsequent spread to other continents. The number of strains encompassed in each pie-chart is indicated as n.

### Population structure of *Clonostachys rosea*

Principal component analysis (PCA) and model-based clustering were performed to investigate population genetic structure among 63 *C. rosea* strains (Table 1 in Supplemental File S1). After LD pruning, 31132 unlinked SNPs were retained for the analysis. The first two principal components, together explaining 38.4% of the genetic variation, separated the strains into six distinct clusters, with the eleven homothallic strains forming a cohesive group (Figure 5A). This pattern was corroborated by the model-based clustering method sNMF, which identified *K* = 6 populations based on the minimum cross-entropy criterion. The eleven homothallic strains were assigned to a single population, whereas the 52 heterothallic strains were distributed across five populations (Figure 5B). Heterothallic populations were mainly structured by geography, with some evidence of admixture. European strains predominated in populations 1, 3, and 4, alongside a minority from America and Canada. Most Chinese strains clustered in population 5, whereas several strains from China, Australia, and America formed population 2 (Figure 5B). Homothallic strains from Central and South America, New Zealand, China, and Japan comprised population 6 (Figure 5B). A neighbour-joining phylogenetic network supported these patterns and revealed multiple reticulations among strains, suggesting shared polymorphisms across lineages (Figure 5C).

**Figure 5.**
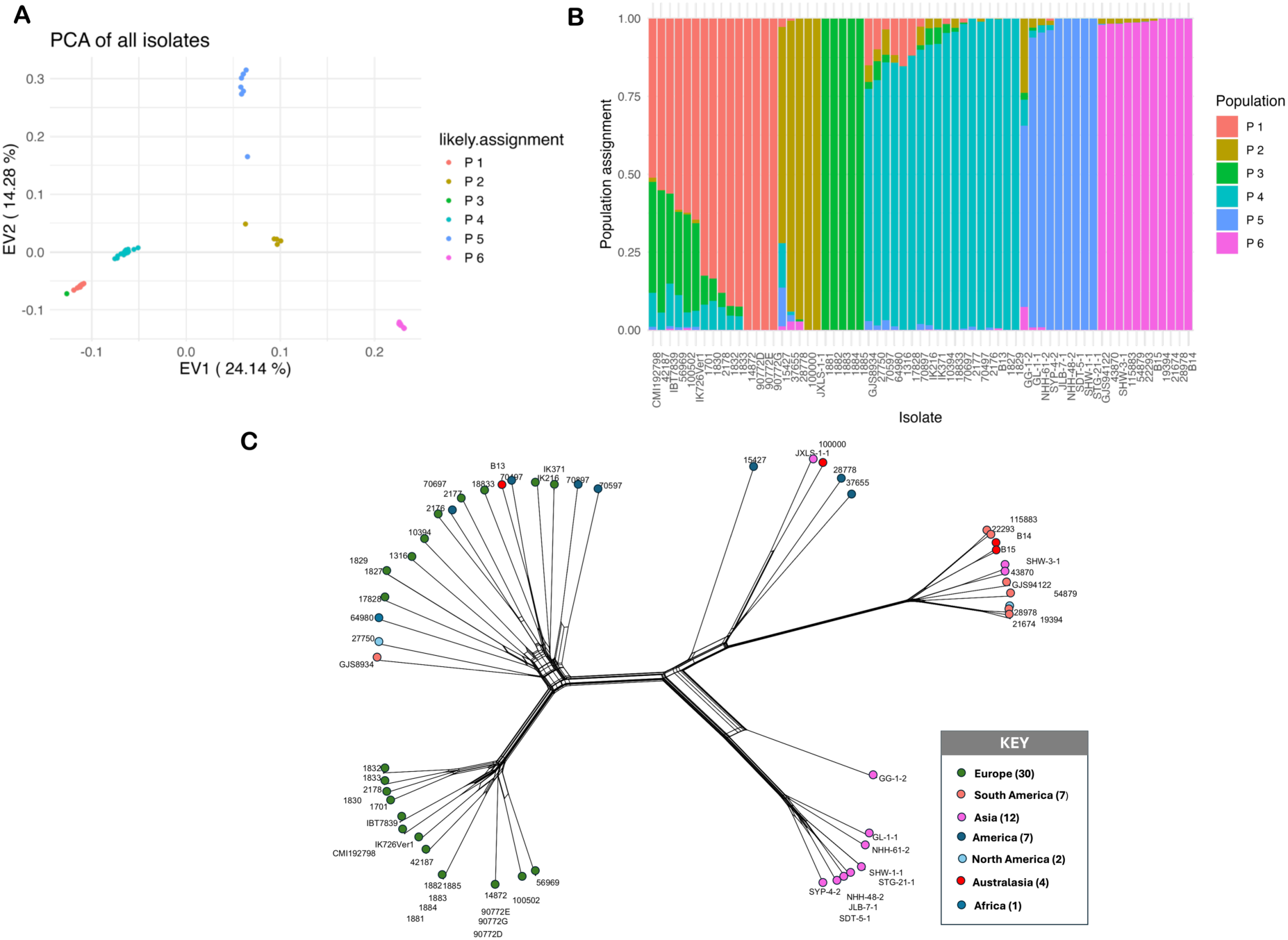
Population structure analyses of heterothallic and homothallic *C. rosea* strains. (**A**) PCA of population structure. Eigenvector 1 (24.14% of variance) is plotted on the x axis and eigenvector 2 (14.28% of variance) is plotted on the y axis. Points are coloured according to the population to which the corresponding strains have been assigned. (**B**) Population structure plot of all 63 strains, with each vertical bar representing a strain coloured according to the population to which it has been assigned. (**C**) Phylogenetic network showing the relationship among all 63 strains. Strain labels are colour-coded according to their region of origin.

### Genetic diversity of *Clonostachys rosea*

Genetic diversity was assessed using minor allele frequency spectra, Tajima’s D, and nucleotide diversity (π) (Figure 6). A total of 1018039 biallelic SNPs were identified across the 63 *C. rosea* strains. After partitioning the whole-genome SNP dataset by mating type (homothallic versus heterothallic) and applying quality filters, 343313 high-confidence SNPs were retained for the homothallic group (11 strains) and 483898 SNPs for the heterothallic group (52 strains).

**Figure 6.**
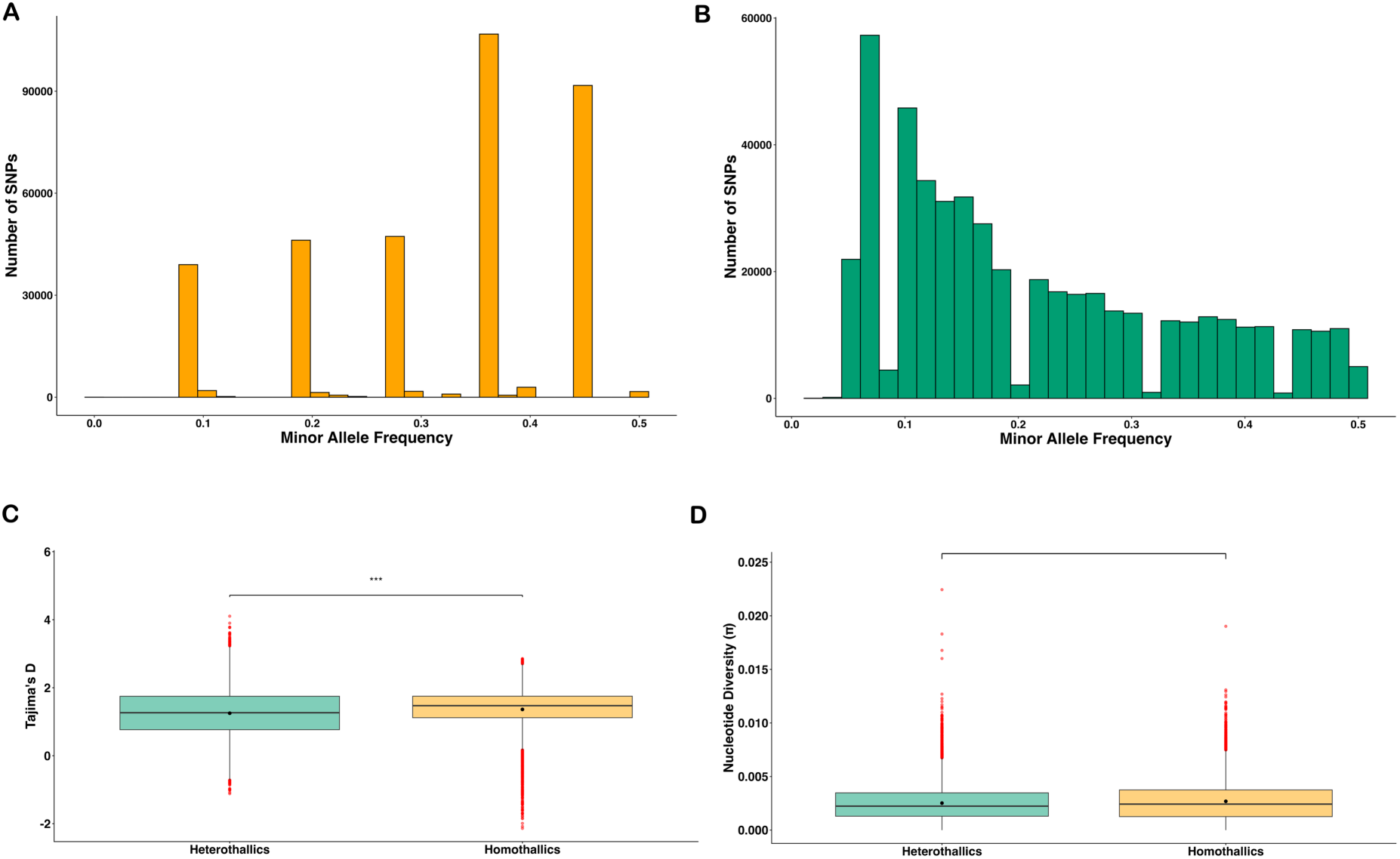
Patterns of genetic diversity in homothallic and heterothallic *C. rosea* strains. (**A**) Minor allele frequency spectrum for SNPs among homothallic *C. rosea* strains. (**B**) Minor allele frequency spectrum for SNPs among heterothallic *C. rosea* strains. (**C**) Comparison of genome-wide Tajima’s D in homothallic and heterothallic groups. Homothallic strains are coloured in orange and heterothallic strains are coloured in green. (**D**) Comparison of genome-wide nucleotide diversity (π) in homothallic and heterothallic groups. Homothallic strains are coloured in orange and heterothallic strains are coloured in green.

In the homothallic group, the minor allele frequency spectrum was enriched for intermediate-frequency variants, with relatively few low- or high-frequency SNPs (Figure 6A). In contrast, the heterothallic group exhibited a left-skewed distribution dominated by low-frequency variants (Figure 6B). Tajima’s D was positive in both groups (homothallic mean D = 1.36; heterothallic mean D = 1.25), indicating an excess of intermediate-frequency SNPs (Figure 6C). A significant difference in mean Tajima’s D between groups was supported by bootstrap resampling (*P* < 0.05).

Despite low overall genetic diversity in both groups, homothallic strains showed slightly higher mean genome-wide nucleotide diversity (π = 0.00269) than heterothallic strains (π = 0.00252); however, this difference was not statistically significant based on bootstrap resampling (*P* > 0.05).

To further characterize recombination history, linkage disequilibrium (LD) decay was estimated separately for each group. LD declined to half of its maximum value at 1150 bp in homothallic strains and 1050 bp in heterothallic strains (Figure 7).

**Figure 7.**
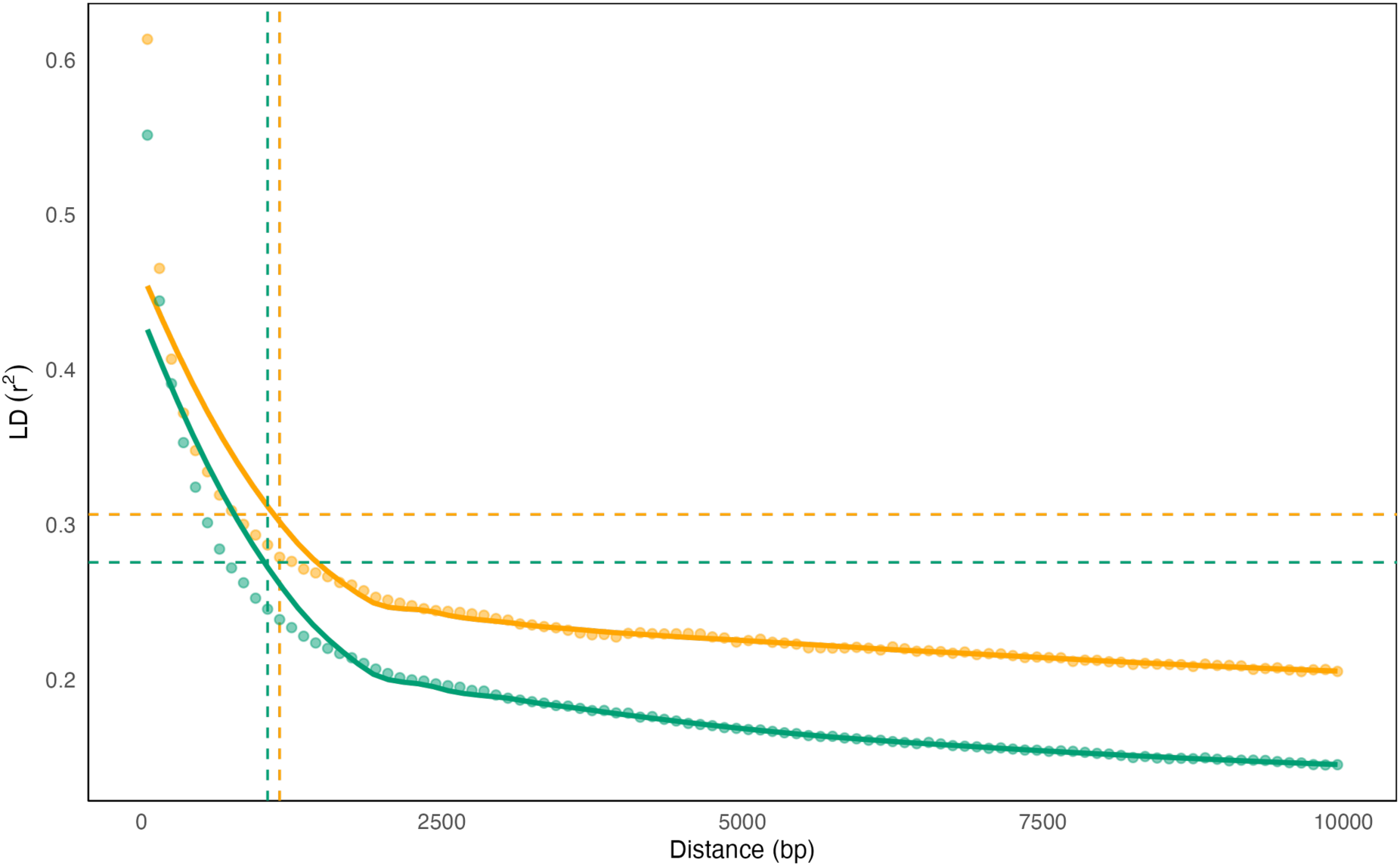
Decay of linkage disequilibrium in homothallic and heterothallic *C. rosea* strains. Linkage disequilibrium (LD) was calculated between pairs of polymorphic sites as a function of the distance between sites in homothallic group, coloured in orange, and heterothallic group, coloured in green. Each point represents the average r^2^ between all pairs of points calculated within sliding 10 kb windows with a step size of 5 kb across the genome. The solid lines are LOESS curves fit to the mean r^2^ values. The vertical dashed lines indicate the position at which LD decayed to half its maximum value in each group. The horizontal dashed lines represent half the maximum LD for each group.

Minor allele frequency distributions were examined for SNPs shared between groups and for SNPs private to each group (Figure 8). SNPs shared between homothallic and heterothallic groups peaked at intermediate allele frequencies (0.2–0.3; Figure 8C), with fewer SNPs at low or high frequencies. Homothallic-specific SNPs were enriched for intermediate-frequency variants, with a peak between 0.3 and 0.4 (Figure 8B), whereas heterothallic-specific SNPs exhibited a left-skewed spectrum dominated by low-frequency variants (Figure 8D).

**Figure 8.**
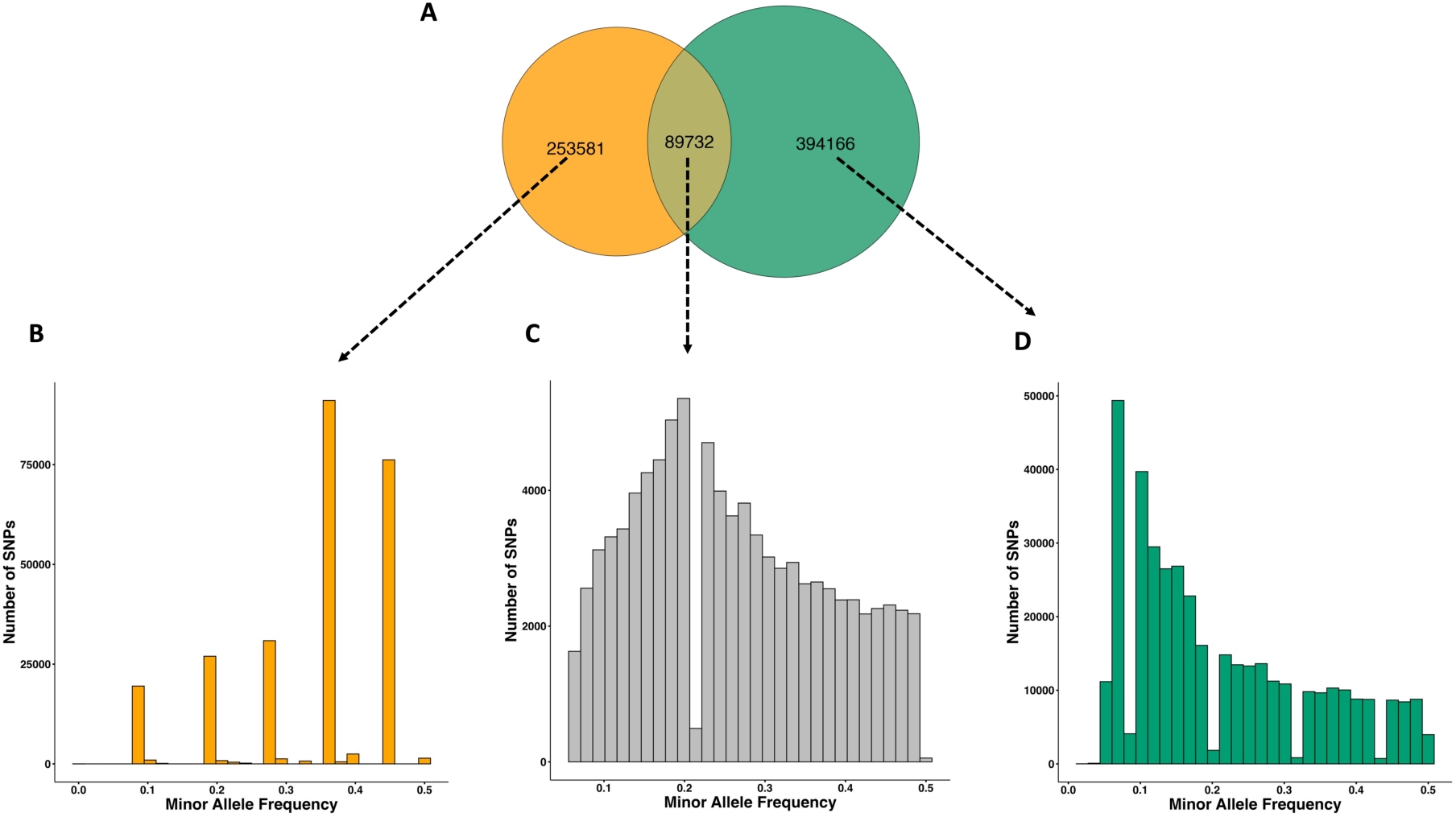
Venn diagram illustrating SNP overlap in homothallic and heterothallic *C. rosea* strains and their corresponding allele frequency spectra. (**A**) Venn diagram showing the overlap and counts of SNPs in homothallic and heterothallic groups. Minor allele frequency spectrum for; (**B**) the SNPs specific to the homothallic group. (**C**) the shared SNPs between the homothallic and the heterothallic groups. (**D**) the SNPs specific to the heterothallic group.

To assess the distribution of functional genetic variation, nucleotide diversity (π) was compared among predicted functional SNP categories (nonsense, missense, synonymous) within shared and group-specific SNP sets (Figure 8). For predicted nonsense SNPs, nucleotide diversity was highest in the shared SNPs (π = 0.0114), intermediate in homothallic-specific SNPs (π = 0.0107), and lowest in heterothallic-specific SNPs (π = 0.0049). Pairwise bootstrap comparisons showed that diversity was significantly higher in homothallic-specific than in heterothallic-specific SNPs (*P* < 0.05), higher in shared than in heterothallic-specific SNPs (*P* < 0.05), and significantly higher in shared than in homothallic-specific SNPs (*P* < 0.05). For predicted missense SNPs, nucleotide diversity was highest in homothallic-specific SNPs (π = 0.0079), followed by shared (π = 0.0064), and heterothallic-specific SNPs (π = 0.0046). Homothallic-specific SNPs showed significantly higher diversity than heterothallic-specific SNPs (P < 0.05), whereas homothallic-specific and shared SNPs did not differ significantly (P > 0.05). Shared SNPs exhibited significantly higher diversity than heterothallic-specific SNPs (P < 0.05). A similar pattern was observed for predicted synonymous SNPs, with the highest diversity in homothallic-specific SNPs (π = 0.0076), followed by shared (π = 0.0064) and heterothallic-specific SNPs (π = 0.0048). Homothallic-specific and heterothallic-specific SNPs differed significantly (P < 0.05), whereas homothallic-specific and shared SNPs did not (P > 0.05); shared SNPs again showed significantly higher diversity than heterothallic-specific SNPs (P < 0.05).

Finally, genome-wide pN/pS ratios calculated across all SNPs differed significantly between groups. The homothallic group exhibited lower pN/pS (0.23) than the heterothallic group (0.29) (Welch’s t-test *P* < 0.001; Wilcoxon test *P* < 0.001), indicating a lower relative proportion of nonsynonymous to synonymous variants in the homothallic lineage.

## Discussion

We investigated the sexual reproductive mode of *C. rosea* to resolve the apparent contradiction between earlier reports of homothallism (Schroers, 1999) and evidence of rapid genome-wide LD decay that typically indicates outcrossing (Broberg et al., 2018). This was achieved through the generation of a population genomic dataset for *C. rosea,* representing its global geographic distribution. Genomic analyses identified both homothallic and heterothallic strains based on the structure and gene content of the *MAT1* locus, and phylogenomics supported a single origin of homothallism within *C. rosea*. Population genomic analyses further showed that homothallic and heterothallic strains formed distinct genetic clusters and differ in allele frequency spectra, Tajima’s D, and LD decay. Together, these results reveal a rare situation in which homothallic and heterothallic lineages coexist within a single fungal species, providing an exceptional opportunity to investigate the genomic consequences of selfing and outcrossing within a shared genomic background.

The 53 to 62 Mb size range of the generated Illumina-based *C. rosea* genome assemblies is consistent with previously reported Illumina-based assemblies, including 58.3 Mb for *C. rosea* IK726 ver. 1 (Karlsson et al., 2015), 56.9 Mb for *C. rosea* ACM941, and 55.5 Mb for *C. rosea* 88-710 (Demissie et al., 2021). Based on the assumption of homothallism, we expected all *C. rosea* genomes to carry both *MAT1-1* and *MAT1-2* idiomorphs. Instead, most strains contained only one of the two *MAT1* idiomorphs, which typically indicates a heterothallic reproductive mode. Presence of the *mat1-2-1* gene was previously reported in *C. rosea* strain IK726 (Karlsson et al., 2015) and is confirmed here. The observation that all strains representing other *Clonostachys* species only carried either *MAT1-1* or *MAT1-2* idiomorphs, but not both, indicates that heterothallism is the ancestral state for the genus, with homothallism representing a relatively recently derived condition in *C. rosea*. Similar phylogenetic patterns of heterothallism as ancestral and homothallism as derived have been reported in multiple ascomycete lineages, including *Botryosphaeriaceae* (Nagel et al., 2018), *Neurospora* (Gioti et al., 2012), and *Cochliobolus* (Yun et al., 1999). The co-occurrence of *MAT1-1* and *MAT1-2* strains in North America, Europe and China, together with short LD decay, suggests the potential for frequent sexual outcrossing and recombination in heterothallic *C. rosea* strains, similar to patterns seen in highly recombining fungal pathogens such as *Zymoseptoria tritici*, where rapid LD decay and balanced mating types ratios indicate regular sexual reproduction (Croll & McDonald, 2012; Feurtey et al., 2023). Earlier work documenting substantial local multilocus diversity of *C. rosea* in a single Danish field is consistent with this picture of historically high gene flow and recombination (Bulat et al., 2000). Heterothallic strains also displayed population genetic structure at the global scale. While this is partly due to geography, where for example, most strains from China are genetically similar, there are also cases where strains from three different continents cluster in the same genetic population, a pattern broadly comparable to other cosmopolitan plant-associated fungi, including *Z. tritici* and *Fusarium graminearum* (Croll & McDonald, 2012; Feurtey et al., 2023; O’Donnell et al., 2000; Wang et al., 2011). The factors driving this population differentiation are currently unknown and will require additional exploration.

Eleven *C. rosea* strains harboured genes from both *MAT1-1* and *MAT1-2* idiomorphs, consistent with homothallism. The localisation of *MAT1-1* and *MAT1-2* genes in tandem on the same contig in at least two strains suggest that the evolutionary mechanism that allowed for the transition from heterothallism to homothallism is an unequal crossing-over event that brought together all genes required for selfing. The tandem localization of *MAT1* idiomorphs due to unequal crossing-over events have been reported from several fungi, including *N. pannonica* and *N. terricola* (Gioti et al., 2012), *F. graminearum* (O’Donnell, 2004), and *Cochliobolus luttrellii* (Yun et al., 1999), suggesting that is a common mechanism for transitions from heterothallism to homothallism. Several homothallic strains produced perithecia in single culture, with asci containing ascospores also being observed. This indicates that the genetic structure of the *MAT1* locus does likely confer homothallism. *Clonostachys rosea* strains originating from ascospores have been reported to frequently produce perithecia in single cultures (Schroers, 1999). Notably, all strains originally reported as the *B. ochroleuca* teleomorph in our current investigation possessed the homothallic version of the *MAT1* locus.

The phylogenomic analysis of the *C. rosea* strains showed that all homothallic strains form a well-supported monophyletic lineage, a pattern reinforced by population structure analyses. This indicates that all homothallic strains included in the current study are descendants of a single unequal crossing-over event that occurred during sexual reproduction between two distinct heterothallic ancestors. The lack of observed anastomosis between homothallic strains indicates that they have accumulated enough genetic variation to be recognised as non-self in vegetative interactions. We hypothesize that this transition occurred in South America, where most homothallic strains were collected. This is further supported by the phylogenetic structure among homothallic strains where a clear distinction can be seen between strains from the south versus the north parts of South America. Under this scenario, the homothallic linage may have first spread within South America and subsequently into Central America and the Caribbean. Furthermore, this scenario suggests at least two independent events of inter-continental movement of homothallic *C. rosea*: one from southern South America to New Zealand, and another from northern South America/Central America to Japan and China. It is tempting to speculate that these intercontinental movements have anthropogenic causes. Comparable long-distance dispersal events have been inferred for other plant- and soil-associated fungi, including *Z*. *tritici* and *Heterobasidion annosum,* where population genomic data implicate anthropogenic drivers such as movement of infected plant material or soil (Croll & McDonald, 2012; Dalman et al., 2010; Feurtey et al., 2023). Given the plant endophytic lifestyle of *C. rosea* (García et al., 2003; Mueller & Sinclair, 1986; Nobre et al., 2005; Walker & Maude, 1975), the movement of plants with associated soil provides a plausible mechanism for the inter-continental spread of the homothallic lineage.

This successful geographic expansion suggests that the homothallic lineage has high ecological fitness despite the potential long-term costs of selfing, although the exact nature of this fitness benefit is currently not known. We can interpret this as an example where homothallism allowed the conservation of a specific combination of beneficial alleles providing high fitness, as proposed for other selfing or weakly recombining lineages of plant-associates, such as *Phytophthora infestans* and mutualists such as *Epichloe* endophytes, where particular genotypes remain successful despite reduced recombination (Drenth et al., 2019; López-Villavicencio et al., 2013; McDonald & Linde, 2002; Schardl et al., 2013). At the same time, comparative analyses of fungal mating systems indicate that transitions from heterothallism to homothallism are often unidirectional, with selfing lineages frequently viewed as evolutionary “dead ends” that rarely revert to outcrossing (Gioti et al., 2012). Classical population-genetic theory further predicts that obligate selfing and the absence of outcrossing promote the accumulation of slightly deleterious mutations through Muller’s ratchet, ultimately reducing fitness and increasing extinction risk (Felsenstein, 1974; Glémin & Galtier, 2012; Muller, 1932, 1964). The coexistence of homothallic and heterothallic lineages in *C. rosea* therefore provides a particularly interesting case to study how the short-term benefits and long-term risks of selfing are manifested at the genomic level.

The structure of the *MAT1* locus itself indeed suggests accumulation of mutations in homothallic strains. This is evidenced by the highly conserved *MAT1* idiomorphs in heterothallic strains compared with the extensive AT-rich repetitive DNA in the homothallic *MAT1* loci. These repetitive regions were sufficiently abundant to prevent full-length assembly of the *MAT1* locus in many homothallic strains, resulting instead in *mat* genes being located on short, fragmented contigs. Similar difficulties in assembling *MAT1* have been reported in other fungi, such as *Sclerotinia borealis*, where repeat-rich *MAT1* regions complicate reconstruction of idiomorph structure (Wilson et al., 2023). In homothallic *Neurospora* species, *mat* genes are reported to evolve under relaxed constraint and to accumulate frameshift and stop codon mutations (Wik et al., 2008). In our experiments, several homothallic *C. rosea* strains produced perithecia, as well as asci with ascospores, despite extensive amounts of repetitive DNA in their *MAT1* locus. These observations show that the *mat* genes are still functional in these strains.

On a genome-wide level, homothallic strains consistently showed higher nucleotide diversity than heterothallic strains within the predicted nonsense, missense and synonymous SNP classes when considering lineage-specific polymorphisms. This pattern is consistent with mutation accumulation under reduced effective recombination (Glémin & Galtier, 2012), although there are also alternative explanations to consider. First, the high nucleotide diversity for nonsense SNPs in shared alleles likely reflects long-standing ancestral polymorphisms that arose before the split between lineages and have been maintained at intermediate frequencies, as observed in other fungi where balancing selection or historical population structure preserves variants across lineages (Feurtey et al., 2023). Second, lower nucleotide diversity in heterothallic-specific SNPs across functional categories is consistent with more efficient purging of new deleterious variants in the recombining lineages, but may also reflect a large pool of recent, low-frequency mutations in the more extensive and structured heterothallic metapopulation. Distinct demographic histories such as bottlenecks or range expansion are known to reduce diversity and generate an excess of rare variants (Charlesworth, 2001), and have been documented in filamentous fungal pathogens with a rapid global spread (Talas & McDonald, 2015).

The lower genome-wide pN/pS ratios observed in homothallic strains, compared to heterothallic ones, is consistent with expectations under the nearly neutral model of molecular evolution (Charlesworth & Eyre-Walker, 2008; Eyre-Walker & Keightley, 2007). Because our estimates are based on standing polymorphism, pN/pS is sensitive to the allele-frequency spectrum and demographic structure. The heterothallic lineage forms a large, globally distributed and structured metapopulation enriched for low-frequency, mildly deleterious nonsynonymous variants, which under nearly neutral dynamics inflate non-synonymous relative to synonymous polymorphism without substantially increasing nucleotide diversity. This provides a parsimonious explanation for the elevated pN/pS in heterothallic strains. In contrast, the homothallic lineage appears recently derived from a recombining ancestor, retains substantial shared ancestral polymorphism, and shows fewer rare variants, consistent with limited time for slightly deleterious non-synonymous mutations to accumulate. Similar time-dependent patterns following transitions to selfing have been reported in recently derived plant selfers such as *Capsella rubella* (Brandvain et al., 2013; Slotte et al., 2013).

A critical question for the evolutionary fate of the homothallic *C. rosea* lineage is whether it is obligately selfing or it occasionally outcrosses, i.e. exhibits facultative homothallism. The elevated Tajima’s D and enrichment for intermediate-frequency alleles in homothallic strains are consistent with limited recombination (Charlesworth, 2001; Tajima, 1989), whereas the left-skewed allele frequency spectrum indicative of an enrichment for low-frequency variants in heterothallic strains is typical of recombining populations (Croll & McDonald, 2012; Feurtey et al., 2023). The intermediate frequencies of shared SNPs suggests that many of these variants were already segregating in the ancestral heterothallic population before the origin of homothallism, similar to patterns inferred in recent transitions to selfing in *C. rubella* (Brandvain et al., 2013; Slotte et al., 2013). The distinct skew towards intermediate-frequency alleles in homothallic-specific SNPs is compatible with limited recombination within the homothallic lineage and suggests that some variants may partly trace back to the two founding individuals that generated this lineage, as expected under recent selfing and founder events where much of the standing variation traces back to a small number of ancestral genotypes (Brandvain et al., 2013; Charlesworth, 2003; Wakeley, 2001).

A well-established measure of the amount of recombination in a population is LD decay, the physical distance over which LD falls to half of its maximum value because of recombination breaking down allele associations (Slatkin, 2008). The similarly short LD decay distances in both homothallic (1150 bp) and heterothallic (1050 bp) lineages indicate extensive historical recombination, comparable to those reported for highly recombining fungi such as *F. graminearum* and *Z. tritici*, where genome-wide LD typically decays within approximately 164 bp to 2 kb (Croll & McDonald, 2012; Feurtey et al., 2023; Singh & Croll, 2021; Talas & McDonald, 2015). The marginally longer LD decay in the homothallic lineage is consistent with a modest reduction in effective recombination under selfing, but we interpret this cautiously because LD is also influenced by effective population size and demographic history (Charlesworth, 2001; Glémin & Galtier, 2012). This pattern has two important implications. First, it suggests either occasional or low-frequency outcrossing in the homothallic lineage, as documented for other homothallic fungi such as *A. nidulans* (López-Villavicencio et al., 2013), or that substantial recombination occurred in the ancestral heterothallic population prior to the origin of homothallism and continues to shape LD patterns. Second, the relatively small difference in LD decay between homothallic and heterothallic strains is consistent with a recent origin of homothallism, such that the full genomic signature of reduced recombination has not yet accumulated. This agrees with theoretical and empirical work showing that genomic signatures of maladaptation in selfing lineages can be subtle and are easily obscured by even low levels of residual outcrossing or by recent transitions to selfing (Escobar et al., 2010; Glémin & Galtier, 2012; Loewe & Cutter, 2008).

The question of occasional outcrossing is also relevant to whether the homothallic lineage should be regarded as a separate species. Under lineage-based species concepts, species are independently evolving lineages (de Queiroz, 1998), and an obligate homothallic lineage could in principle, be viewed as a separate species. In *C. rosea*, the homothallic lineage is clearly divergent in genome-wide phylogenies and mating-type organisation, yet we still observe shared polymorphisms, low but detectable admixture, and reticulation in the neighbour-net. These observations suggest that complete reproductive isolation has not yet evolved. On this basis, we therefore view the homothallic lineage as a strongly differentiated, relatively recently derived lineage that is still undergoing divergence from heterothallic *C. rosea*, rather than a fully separated species.

Resolving the genetic population structure of *C. rosea* has important applied implications. Population genomic datasets have been used extensively to dissect agriculturally important traits in fungi through genome-wide association (GWAS) mapping, including virulence in pathogens such as *H. annosum* (Dalman et al., 2013), *Z. tritici* (Gao et al., 2016), *Rhynchosporium commune* (Mohd-Assaad et al., 2016), and *F. graminearum* (Talas & McDonald, 2015). As some *C. rosea* strains are used commercially as biological control agents (Jensen et al., 2022), similar approaches could be used to identify genetic determinants of high biocontrol efficacy, which in turn can be used as markers for improving biocontrol agents. However, including homothallic strains from a largely selfing lineage in association panels that otherwise consist of recombining heterothallic strains may introduce artefacts by violating GWAS assumptions regarding recombination and linkage structure (Croll & McDonald, 2012). Previously, Broberg et al. (2018) reported a *C. rosea* GWAS of growth rate at cold temperature and Iqbal et al. (2020) reported a similar study on the ability of *C. rosea* to produce nematicidal compounds. However, only a single homothallic strain (CBS 193.94) was included in these studies, which is unlikely to have strongly biased those results. Based on the current work, we recommend that future GWAS exclude homothallic strains.

In summary, we generated a population genomic dataset for *C. rosea* and identified a unique combination of homothallic and heterothallic lineages within a single fungal species. By integrating *MAT1* locus structure, phylogenomics, population genetic metrics (allele-frequency spectra, nucleotide diversity, Tajima’s D, LD decay), and pN/pS, we reconcile the apparent discrepancy between previously reported homothallism and rapid LD decay and infer a recent origin and successful geographic expansion of a homothallic lineage. Our data provide important practical guidance for future population genomic studies aimed at identifying the genetic basis for biocontrol-related traits in *C. rosea*.

## Author contributions

DMW, AW, MD, MBD and MK planned and designed the experiments. BJ, AR and MD performed the lab work, while DMW, AW, EP, MBD, MB, SC, HDFL, MD and MK analysed the data. All authors read and approved the manuscript.

## Acknowledgements

This study was financially supported by the Swedish Research Council for Environment, Agricultural Sciences and Spatial Planning (FORMAS) (grant number 942-2015-1128), by the Department of Forest Mycology and Plant Pathology at the Swedish University of Agricultural Sciences, and by SLU Grogrund. Sequencing was performed by the SNP&SEQ Technology Platform in Uppsala. The facility is part of the National Genomics Infrastructure (NGI) Sweden and Science for Life Laboratory. The SNP&SEQ Platform is also supported by the Swedish Research Council and the Knut and Alice Wallenberg Foundation.

## Data availability

Sequencing reads and annotated genomes are available on European Nucleotide Archive (ENA) under bioproject PRJEB108086.

## Supplementary material

Supplemental File S1, Table 1. *Clonostachys rosea* strains used in the current study.

Supplemental File S1, Table 2. RNA-seq datasets used for gene prediction in the current study.

Supplemental File S1, Table 3. Genomes from additional species and strains used in the current study.

Supplemental File S1, Table 4. Genome statistics for *Clonostachys rosea* genomes generated in the current study.

Supplemental File S1, Table 5. Distribution of mating type idiomorphs in *Clonostachys* strains.

## Notes

### Competing Interest Statement

The authors have declared no competing interest.

